# Deep Geometric Framework to Predict Antibody-Antigen Binding Affinity

**DOI:** 10.1101/2024.06.09.598103

**Authors:** Nuwan Bandara, Dasun Premathilaka, Sachini Chandanayake, Sahan Hettiarachchi, Vithurshan Varenthirarajah, Aravinda Munasinghe, Kaushalya Madhawa, Subodha Charles

## Abstract

In drug development, the efficacy of an antibody depends on how the antibody interacts with the target antigen. The strength of these interactions gives an indication of how successful an antibody is in neutralizing an antigen. Therefore, the strength, measured by “binding affinity”, is a critical aspect of antibody engineering. In theory, the higher the binding affinity, the higher the chances are that the antibody is successful against the target antigen. Currently, techniques such as molecular docking and molecular dynamics are utilized in quantifying the binding affinity. However, owing to the computational complexity of the aforementioned techniques, running simulations for large antibodies/antigens remains a daunting task. Despite the commendable improvements in deep learning-based binding affinity prediction, such approaches are highly dependent on the quality of the antibody-antigen structures and they tend to overlook the importance of capturing the evolutionary details of proteins upon mutation. Further, most of the existing datasets for the task only include antibody-antigen pairs related to one antigen variant and, thus, are not suitable for developing comprehensive data-driven approaches. To circumvent the said complexities, we first curate the largest and most generalized datasets for antibody-antigen binding affinity prediction, consisting of both protein sequences and structures. Subsequently, we propose a deep geometric neural network comprising a structure-based model and a sequence-based model that considers both atomistic and evolutionary details when predicting the binding affinity. The proposed framework exhibited a 10% improvement in mean absolute error compared to the state-of-the-art models while showing a strong correlation between the predictions and target values. We release the datasets and code publicly (https://drug-discovery-entc.github.io/p2pxml/) to support the development of antibody-antigen binding affinity prediction frameworks for the benefit of science and society.

## Introduction

In the field of drug development, molecules can be categorized as large and small molecules, based not only on their size but also on how they are synthesized, mode of action, transportation, etc. Small molecules are stable, synthetic chemicals with relatively simple structures. These have been around for decades and constitute the majority of the drugs currently in use. The body can absorb small molecules through oral uptake. Common small molecule drugs include “medicine cabinet” drugs such as aspirin and penicillin. Large molecules, also known as “biologics”, have complex structures and majorly consist of proteins produced by living cells. Their production processes are complicated and time-consuming. Examples of biologics in use include vaccines, blood/blood components, etc., and these are usually administered through injections. Biologics have high specificity in destination targeting, whereas small molecules may bind to off-targets and induce non-target harmful effects (i.e. side effects). Eight out of ten global best-selling drugs being biologics in 2018, indicates the increasing significance of biologics in the field of pharmaceuticals (1).

The efficacy of drugs designed using biologics (and even using small molecules) depends on how well they can bind or interact with the target molecule(s). Thus, the strengths of those interactions must be evaluated during the drug design phase to achieve the desired efficacy levels. The *binding affinity* reflects the strength of the interactions. The free energy associated with the binding of two molecules is called the binding free energy. Binding energy, which is generally a negative figure, is taken as a quantitative measure of the binding affinity. Accordingly, the higher the binding energy, the higher will be the binding affinity. Therefore, accurate binding energy prediction is a helpful tool for designing drugs with higher affinity and specificity towards their target.

In this work, we focus on antibody-antigen binding, which is a type of biologics interaction. Antigens are foreign substances that induce an immune response in a body, whereas antibodies are a part of the immune response produced to fight off such antigens. The extent of interactions between an antibody-antigen pair, evaluated through the binding affinity, decides the suitability of the drug in subsiding the relevant pathogenic condition. Here, we primarily consider the IC50 value, which is the concentration of the antibody required to inhibit the antigen activity by 50%, as the quantitative measure for the binding affinity. Further, we only focus on antibodies and antigens that are classified as proteins in this work.

Currently, molecular docking is used to study how two or more molecular structures fit together (2) through computer simulations. However, modeling interactions between two molecules is complicated as this involves a range of forces. To produce a stable bond between molecular structures, the binding naturally happens via the lowest energy pathway. Molecular docking aims to mimic these natural interactions that take place during the binding via the lowest energy pathway. To achieve this, molecular docking calculates binding affinities at different binding poses to determine the optimum binding pose and binding affinity. However, the accuracy of molecular docking heavily depends on the ability of the utilized potential function to describe the forces in the system in a precise manner. In contrast, molecular dynamics (MD) simulation is a more sophisticated simulation technique that extends the capabilities of molecular docking by including the temporal behaviour of the structures of concern. Due to the over-dependency of MD simulations on the number of atoms in the proteins of concern, conducting MD simulations for large molecules remains a daunting task even to this date. To this end, recent advancements in deep learning have led researchers to use deep learning models as a faster alternative to time-consuming MD simulations (3, 4). But, the predictive performance of existing deep learning methods when calculating the binding affinity is highly dependent on the quality and resolution of the three-dimensional structures of the antibody and antigen.

To overcome the challenges of the above traditional methods, numerous deep learning-based methods have recently been utilized to predict the binding affinity. To this end, Wang et al. (5) introduced a method involving topology-based feature generation using element and site-specific persistent homology to capture the structural characteristics of 3D protein structures. This method employs a combination of convolutional neural network (CNN) and gradient boosting tree (GBT) model in which the CNN is utilized to extract more concise features from the topological descriptors, followed by the GBT as the prediction model. However, this approach is limited to predicting the effects of mutations within protein-protein complexes. In (6), the authors presented a geometric attention network that generates embeddings for mutation and wildtype 3D complexes, capturing residue information based on atom proximity while the attention is employed to identify crucial residue pairs at the protein interface for binding affinity. However, in this approach, capturing evolutionary details solely through 3D structures is ineffective and presents poor performance. Li et al. (7) proposed a bi-directional attention neural network for predicting compound-protein interactions and binding affinity. This method models compound-protein interactions as a continuous value prediction problem and employs graph neural networks (GNNs) to process structural data, alongside convolutional neural networks (CNNs) for processing sequence data. The network integrates both representations but has limitations in efficiently capturing residue features due to fixed windowing over protein sequences. More recently, DG-affinity (8) proposed to utilize large-language models to process amino-acid sequences to predict binding affinity, which still lacks the capability to capture the atomistic information through the structures for affinity prediction.

In this work, we introduce a novel deep-learning model that combines a geometric model utilizing graph convolution and graph attention operations to process the antibody-antigen structures and a sequence model utilizing self-attention and cross-attention to model the amino-acid sequences of the antibody-antigen pair. One major goal of the study was to develop a general deep model that is not confined to a specific family of antigens. Accordingly, the proposed network was trained on a curated dataset comprising antibody-antigen pairs for HIV, MERS, flu virus, etc. We observed a significant improvement in the mean absolute error compared to existing state-of-the-art models. Therefore, the key contributions of this work can be summarized as follows:

- We curate the largest and most generalized datasets for antibody-antigen binding affinity prediction in the literature, including both protein sequences and structures.
- We propose an end-to-end deep learning framework for antibody-antigen binding affinity prediction that combines a geometric model, which processes the atomistic-level structural details of the antibody-antigen pairs, and a sequence model, which processes the evolutionary details of the antibody-antigen pairs. Through extensive evaluations on the existing datasets and our curated datasets, we show that the proposed framework consistently outperformed the state-of-the-art methods by significant margins.
- We present a web-based platform where the users can obtain predictions for the binding affinity of their desired antibody-antigen pairs via uploading protein structure files. Further, we release the codes and datasets openly to support the development of antibody-antigen prediction frameworks for the benefit of science and society.

## Results

In this section, we discuss in detail our experiments and results in comparison with several baseline models from the literature. The impact of incorporating both evolutionary and atomistic-level details through sequence-based and structure-based models, respectively, is exhibited in the results. The results are presented and discussed in the following order of subsections: Dataset Curation, Combined Models, Sequence-based Models, and Structure-based Models.

### A. Dataset Curation

As publicly available datasets are compiled in different formats, they were initially preprocessed through custom pipelines and the resulting curated sequence and structure datasets were named *P2PXML-Seq* and *P2PXML-PDB*, respectively. A detailed comparison of the curated datasets with the publicly available datasets is provided in Table 1. P2PXML-Seq only contains antibody-antigen pairs in the respective amino acid one-letter code (FASTA format (9)) while the P2PXML-PDB contains 3D structures of the antibody-antigen pairs in protein data bank (PDB) format.

**Table 1.** Dataset comparison. *Here, usable datapoints refer to the remaining datapoints after a defined set of preprocessing steps, including the duplicate removal and the exclusion of datapoints with no numerical value for binding affinity. **P2PXML-Seq and P2PXML-PDB are our curated protein sequence and structure datasets, respectively.

| Dataset | Usable Datapoints* |
| --- | --- |
| AB-Bind (10) | 1 101 |
| Ab-Cov (11) | 1 420 |
| CATNAP (12) | 11 208 |
| AlphaSeq (13) | 87 808 |
| SAbDab (14) | N/A |
| Skempi (15) | N/A |
| P2PXML-Seq (Ours)** | **111 845** |
| P2PXML-PDB (Ours)** | **8 475** |

As per our knowledge, the datasets in Table 1 are the most extensive datasets curated explicitly for the antibody-antigen binding affinity prediction. Our datasets have a consistent format concerning the antibody-antigen pairs and the corresponding numerical values (i.e. IC50; half-maximal inhibitory concentration). It is to be noted that we assumed specific parameters or conditions including non-competitive inhibition, following (16, 17), to approximate IC50 whenever needed. Moreover, the curated datasets express sufficient generalizability since they contain numerous antigens such as SARS-CoV-2, HIV, MERS, and flu, as shown in Figure 1, and their related antibodies, in contrast to many datasets in the literature where only one antigen is considered (11, 13).

**Fig. 1.**
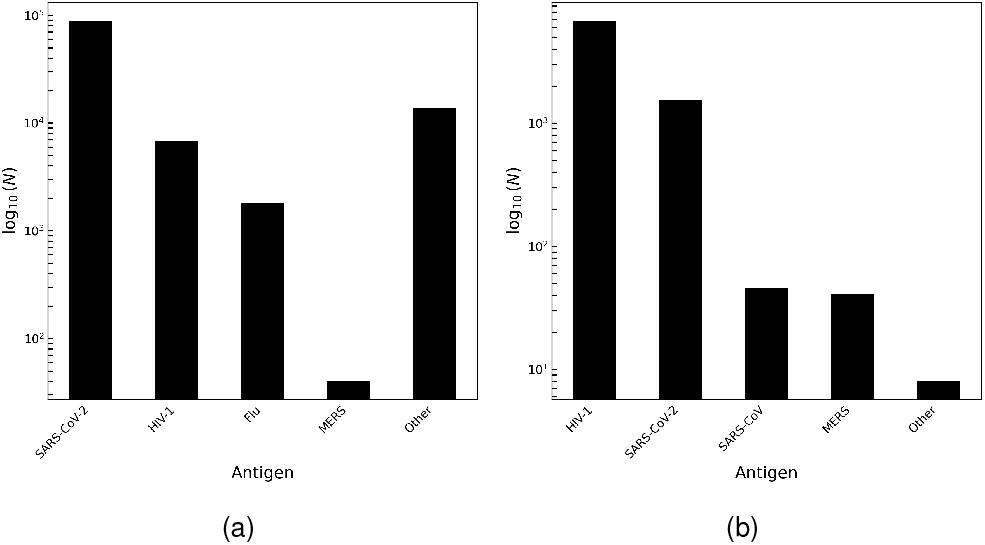
The number of antibody-antigen pairs (in log scale) vs the type of antigen strain in the curated (a) P2PXML-Seq dataset and (b) P2PXML-PDB dataset. Here, *N* refers to the number of antibody-antigen pairs and “Other” refers to antigen strains, which do not correspond to a specific bar in the plots, such as WIV1 (18) and SHC014 (19).

### B. Combined Models

The primary intuition behind the combined model is to incorporate the evolutionary and atomistic-level details of antigens and antibodies through the sequence-based model and the structure-based model, respectively, while sharing the information learned through each pipeline to imitate the chemical binding potential. Further, as depicted in Figure 2, it is evident that antibodies corresponding to a particular antigen variant do not exhibit distinctive clusters in low-dimensional space in both protein sequences and structures which further emphasizes the importance of modeling an information sharing pipeline between antibodies and antigens while utilizing both their sequences and structures.

**Fig. 2.**
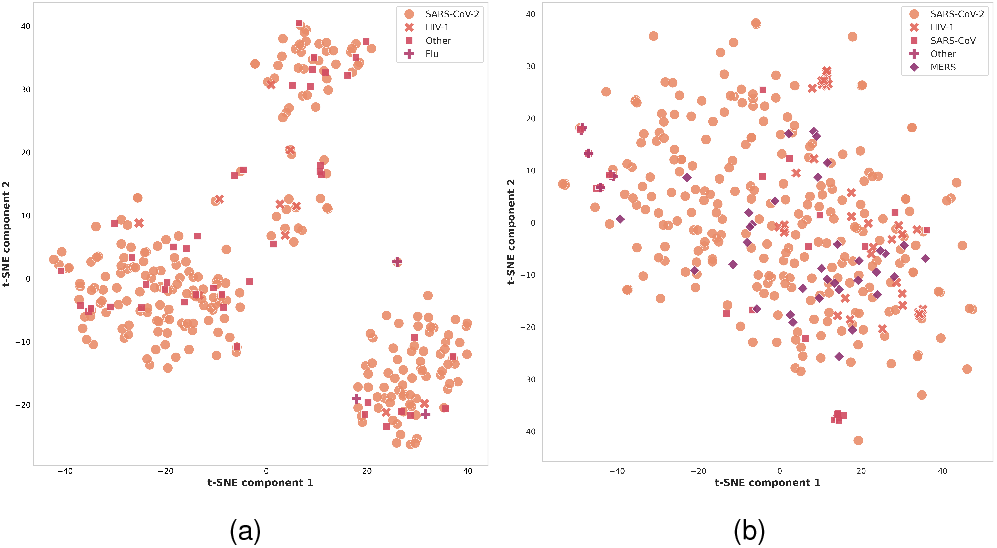
Component 2 vs component 1 of the low-dimensional t-distributed stochastic neighbor embeddings (20) reduced from the (a) amino-acid sequence embeddings generated by following ESM-2 (21). The sequences are from a randomly-sampled subset of the antibody amino-acid sequences in the P2PXML-Seq dataset. (b) whole-graph embeddings of the constructed graphs. The graph embeddings are generated by following graph2vec (22) and the graphs are corresponding to a randomly-sampled subset of the antibody protein structures in the P2PXML-PDB dataset. Here “Other” follows the same definition as in Figure 1.

The curated P2PXML-PDB dataset was used to train, test and compare the performances of the models. As the proposed network comprises a sequence model in conjunction with a structure model, the results exhibited by the proposed network were compared with state-of-the-art sequence-based and structure-based models.

From Table 2, it is evident that our final Combined-V2 model outperforms all the considered state-of-the-art approaches at least by a margin of 10.6% while improving the performance of our individual sequence-based and structure-based models by 5.6% and 6.8%, respectively. The performance of the proposed combined model underscores the importance of incorporating both atomistic-level and evolutionary details in predicting the binding affinity. Furthermore, as depicted in Figure 3, the correlation between the predicted and target values is significantly strong: the Pearson correlation coefficient being 0.8703 and the Spearman’s correlation coefficient being 0.9450 which further highlights the better performance of our approach.

**Table 2.** Overall results comparison for P2PXML-PDB dataset with respect to mean absolute error (MAE) and mean squared error (MSE). Here, ‘1D-CNN with attention’ and ‘GCN’ refer to the respective state-of-the-art sequence and structure models whose performances are compared with our sequence-based pipeline (Parallel Transformer with cross-attention), our structure-based pipeline (Parallel GCN + GATConv) and the final combined model containing both pipelines (Combined-V2 + pre-trained weights). Only the best-performing structure and sequence models out of the considered work found in the literature are given. Other related research is mentioned under the subsections; Sequence-based Models and Structure-based Models.

| Model | MAE | MSE |
| --- | --- | --- |
| 1D CNN with attention (23) | 1.1191 | 5.2266 |
| GCN (24) | 1.1348 | 5.2988 |
| Parallel Transformer with cross-attention (Ours) | 1.1083 | 5.1986 |
| Parallel GCN + GATConv (Ours) | 1.1338 | 5.3457 |
| Combined-V2 + pre-trained weights (Ours) | **1.0005** | **4.6709** |

**Fig. 3.**
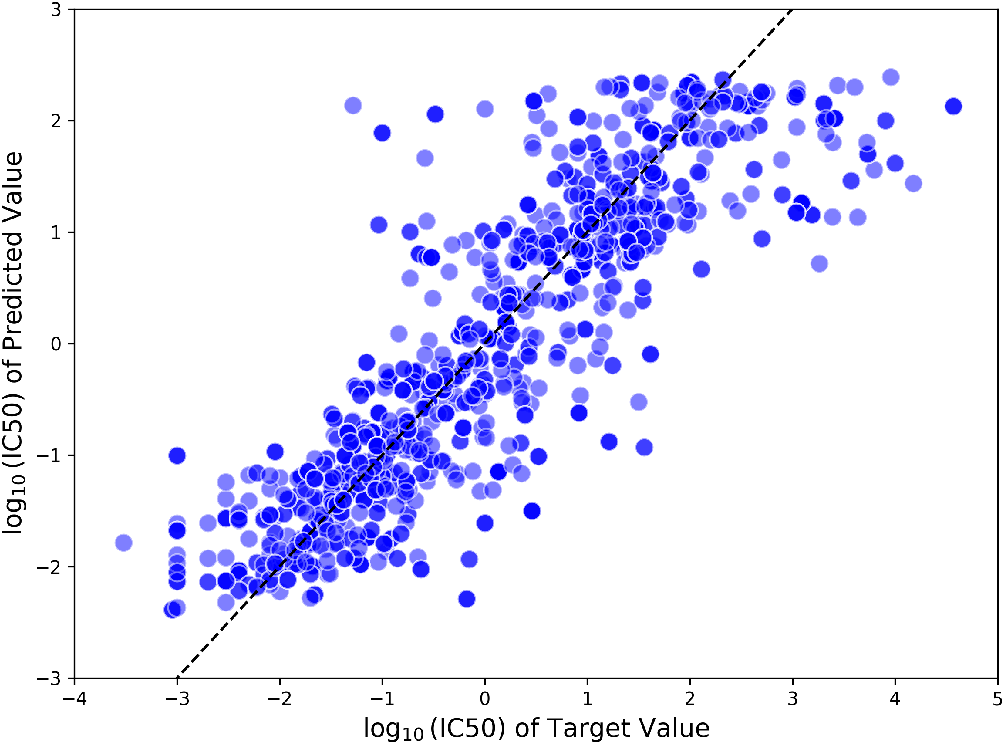
*log*_10_ (*IC*50) of the predicted values vs *log*_10_ (*IC*50) of the target values for the test set of P2PXML-PDB dataset using our best performing model. The Pearson correlation coefficient is 0.8703 and the Spearman’s correlation coefficient is 0.9450 between the predicted and target values.

In our experiments, several variants of the combined model were tested along with two different amino acid encoding schemes. An overview of variations and performance comparison between the combined model variants are given below. See section G for more details.

- Combined-B: This refers to the combined base model where the outputs from the sequence-based model and the structure-based models are collectively considered for calculating the combined loss function.
- Combined-V1: The Combined-B model was modified such that the output latent vectors from sequence-based and structure-based models were concatenated. Then the concatenated vectors were propagated through parallel paths to obtain separate outputs which were eventually combined to calculate the combined loss function.
- Combined-V2: In contrast to the direct concatenation of latent vectors from two pathways as mentioned in the Combined-V1 model, here, the resulting vector from cross-attention between the two pathways was concatenated separately with each latent vector from two models. The resulting concatenated vectors were processed similarly as in Combined-V1. See Figure 7.
- Combined-V2 + pre-trained weights: Instead of randomly initializing the weights of the sequence model in the Combined-V2 model, pre-trained weights obtained from training the sequence model using the P2PXML-Seq dataset were utilized. Note that the weights in the structure model of the Combined-V2 model were still initialized randomly. Since the Combined-V2 model was trained on the P2PXML-PDB dataset, pre-trained weights were not used for the structure-based counterpart in the Combined-V2 model as it would have otherwise pre-exposed the dataset to the model.

As given in Table 3, the Combined-V1 model outperformed the Combined-B model by a margin of 5% which highlights the impact of sharing evolutionary and atomistic details between sequence-based and structure-based pipelines. Applying cross-attention when sharing information between the sequence and structure models in Combined-V2 further improved the performance of Combined-V1, validating the importance of learning the extent to which information should be shared. At last, employing pre-trained weights, as described earlier, slightly improved the results further to obtain the best MAE of 1.0005.

**Table 3.** Results comparison between the principal component score vector of hydrophobic, steric, and electronic properties (VHSE8) encoding and pre-trained protein language embeddings from ProtT5-XL-BFD (25) for protein amino-acid sequences. The dataset used is our P2PXML-PDB dataset and the performance parameter is MAE. *VHSE8 encoding of an amino acid is an 8D Vector of Hydrophobic, Steric, and Electronic properties. **Obtained from ProtT5-XL-BFD which is a pretrained model on protein sequences.

| Model | MAE |  |
| --- | --- | --- |
|  | VHSE8 encoding* | Protein language embeddings** |
| Combined-B | 1.0567 | 1.1209 |
| Combined-V1 | 1.0039 | 1.0988 |
| Combined-V2 | 1.0007 | 1.0951 |
| Combined-V2 + pre-trained weights | **1.0005** | **1.0924** |

It can be deduced from Table 3 that the same pattern of performance improvement was perceived when utilizing the protein language embeddings from ProtT5-XL-BFD (25) instead of the principal component score vector of hydrophobic, steric, and electronic properties (VHSE8) encoding scheme to encode the protein amino-acid sequences. However, the superior performance of VHSE8 encoding over protein language embedding in every combined model variant suggests that it is better to use an encoding scheme rather than deep learning-based protein language embeddings, given the fact that we utilized the language embeddings from a pre-trained model rather than a model which was fine-tuned to the task in hand. We further believe that if we fine-tune the protein language model to our task, it might have a comparable or superior performance than the traditional encoding scheme since it could learn an enriched protein sequence representation tailored to the task rather than an ill-posed generalized representation which would not be sufficient to obtain the best performance.

### C. Sequence-based Models

The sequence-based model was developed to capture evolutionary details of the amino acid sequences of both antigens and antibodies while hierarchically sharing the information learned from parallel pipelines through cross-attention. Based on this intuition, several approaches were tested and then compared with the state-of-the-art approaches (23, 27), which utilize protein sequences for the binding affinity prediction.

As per the results in Table 4, the sequence-based model with the best performance is the ‘Parallel Transformer with cross-attention’ model. The results indicate that the selected sequence-based model surpasses the state-of-the-art approaches under all three datasets by MAE margins of 4%, 44%, and 1% for VirusNet, P2PXML-Seq, and P2PXML-PDB datasets, respectively. Consequently, it was incorporated into the combined model to extract the information from protein amino acid sequences.

**Table 4.** Results comparison between the protein sequence-based models with MAE as the performance parameter. The datasets used are VirusNet (26), P2PXML-Seq and P2PXML-PDB. Apart from ‘SVR with RBF kernel’ and ‘1D CNN with attention’ models, the other models were developed during our study to check the impact of different architectures on the MAE. *Here, ‘Transformer with multiple cross-attention’ refers to having cross-attention blocks in each stage, in addition to the two hierarchical cross-attention layers as in ‘Parallel Transformer with cross-attention’. **In the ‘Transformer with distogram’ model, the calculated distograms were used as input instead of the encoded protein sequences. ***The pre-trained protein language embeddings generated through ProtT5-XL-BFD (25) (without fine-tuning the model to our task) were used as input to the transformer instead of encoded protein sequences.

| Model | Parallel Pipeline | Cross attention | MAE |  |  |
| --- | --- | --- | --- | --- | --- |
|  |  |  | VirusNet (26) | P2PXML-Seq | P2PXML-PDB |
| SVR with RBF kernel (27) | ✗ | ✗ | 102.1267 | 10.8236 | 2.4181 |
| 1D CNN with attention (23) | ✓ | ✓ | 3.5124 | 0.0964 | 1.1190 |
| LSTM | ✓ | ✗ | 5.2853 | 0.2460 | 1.4448 |
| LSTM | ✓ | ✓ | 3.6288 | 0.0926 | 1.1275 |
| Transformer | ✓ | ✗ | 3.4995 | 0.0898 | 1.1175 |
| Parallel Transformer with cross attention | ✓ | ✓ | **3.3707** | **0.0542** | **1.1083** |
| Transformer with multiple cross-attention* | ✓ | ✓ | 3.4744 | 0.0553 | 1.1084 |
| Transformer with distogram** | ✓ | ✓ | 3.3882 | 0.0551 | 1.1108 |
| Transformer with protein language embeddings*** | ✓ | ✓ | 3.4263 | 0.0623 | 1.1128 |

### D. Structure-based Models

The structure-based model was developed to capture atomistic-level and residue-level structural information of the antibodies and antigens to facilitate the antibody-antigen binding affinity prediction. Based on this intuition, several approaches were tested and compared with the state-of-the-art approaches (24, 28), which utilized protein 3D structures for the binding affinity prediction.

As per the results in Table 5, the final structure-based model is selected to be the ‘Parallel Graph Convolution (GCN) + Graph Attention (GATConv)’ model. The results depict that the proposed structure-based model has surpassed the state-of-the-art methods under our benchmark PDB dataset by a slight MAE margin of 0.08% and an MSE margin of 0.04%. Accordingly, it was integrated into the combined model to extract information from protein 3D structures.

**Table 5.** Results comparison between the protein structure-based models with MAE and MSE as the performance parameters. The dataset used is our P2PXML-PDB dataset. *Here, the parallel GAT model refers to a full graph attention network, which was developed under the hypothesis that identifying the most essential nodes through attention would be sufficient for better binding affinity prediction. However, the performance of the model indicates that such information alone is inadequate to yield better performance. **The parallel GCN model with cross-attention was developed following the success of our final sequence-based model with the expectation that the information sharing between the parallel paths would be beneficial for a better prediction. Surprisingly, it was not as useful in the structure-based model as it was in the sequence-based model. We hypothesize that the local aggregation of the node features in the intermediate stages lacks or does not encapsulate sufficient global information, which is essential for the graph-level prediction task.

| Model | MAE | MSE |
| --- | --- | --- |
| Parallel GNN (28) | 1.1633 | 5.5406 |
| Parallel GCN (24) | 1.1347 | 5.3490 |
| Parallel GAT* (Ours) | 1.1788 | 5.6719 |
| Parallel GCN + cross-attention** (Ours) | 1.1409 | 5.3818 |
| Parallel GCN + GATConv (Ours) | **1.1338** | **5.3457** |

## Conclusions

In the field of drug development, the efficacy of a drug depends on the extent to which the constituent molecules interact with the target molecules. Thus, the strengths of those interactions must be evaluated during the drug design phase to achieve the desired efficacy levels. In literature, such protein-protein interactions are reflected by the binding affinity. Therefore, accurate binding affinity prediction is critical in designing drugs with higher specificity towards the target. Our study focused on antibody-antigen binding, which is a subclass of protein-protein interactions. Currently, techniques such as molecular docking and molecular dynamics simulations are employed to determine the binding affinity at different binding poses, but they either overlook the temporal behaviour, leading to lesser accuracy (in Molecular Docking) or are computationally expensive and time-consuming (in Molecular Dynamics). Even though there is a trending interest in utilizing machine learning to predict binding affinity, the predictive performance of existing machine learning methods when calculating the binding affinity is highly dependent on the quality of the antibody-antigen structures, and they tend to overlook the importance of capturing the evolutionary details of proteins upon mutation.

To overcome the said complexities and drawbacks, we proposed a novel deep geometric network that comprises a structure model that could process the 3D structures of the input proteins and a sequence model that could handle the amino acid sequences of the input proteins. We employed attention mechanisms in both models to ensure that both atomistic-level information and evolutionary details are appropriately incorporated into neighbor embeddings and/ or between the pipelines for antibodies and antigens. The proposed model was trained on our curated dataset, which consists of sequences and structures of antigens and antibodies corresponding to a diverse set of common viruses such as Human Immuno-deficiency Virus (HIV), SARS-CoV-2, etc., to ensure sufficient generalizability within the dataset.

After extensive ablation studies which were performed to select the encoding schemes, node and edge features, and model hyperparameters, it was observed that the Combined-V2 model, which is our final model architecture including selected sequence-based and structure-based models, surpassed the state-of-the-art approaches at least by a margin of 10.6% in terms of the mean absolute error. Furthermore, the stand-alone sequence-based model was able to surpass the existing sequence-based state-of-the-art methods under all benchmark datasets, and our structure-based model marginally outperformed the existing works in the literature as well.

Moreover, we developed a website as a community access tool to allow the interested community to obtain prediction results for their input proteins via our hosted models.

In summary, we believe that the work presented in this study would be beneficial in improving deep-learning-based binding affinity prediction of antibody-antigen pairs, especially in the domain of drug development.

## Methods

In this section, we delve into the details of the dataset curation process and our proposed models: sequence model and structure model followed by the model that combined both sequence and structure models. We discuss the mathematical concepts associated with deep-learning models and the justifications for employing specific techniques. In brief, the sequence model processes the amino acid sequences of the input proteins, whereas the structure model treats the 3D structures of the proteins as graphs with nodes and edges. In addition to the sequence and structure models, the combined model has attention layers to share information between the said models.

### A. Dataset Curation

Since publicly available datasets are tabulated in different formats, the first task associated with curating a generalized dataset was to process the datasets to have a similar format. In addition, as the models require the 3D structure of the proteins, suitable measures had to be taken to generate the 3D structures that were unavailable in some public datasets. Homology modeling (29) and AlphaFoldV2 (30) were employed in this regard. The Table 6 presents a summary of publicly available datasets. Generally, experimental uncertainties can produce multiple IC50 values for a given antibody-antigen pair over several trials. To negate the impact of outliers, we first considered the median value of the provided IC50 values for repeated entries of antibody-antigen pairs. Based on the availability of the template structure and the mutation profile, we then decided whether to use Homology Modelling or AlphaFoldV2 to generate the 3D structure. If the mutation profile, along with the template structure, was provided, then we utilized homology modeling. If either of the two requirements was not satisfied, then we used the AlphaFoldV2 pipeline. Figure 4 shows the processing steps associated with the dataset curation.

**Table 6.** Summary of the used publicly available datasets. Here, ΔΔ*G*, IC50, EC50, IC80, ID50 and *K*_*d*_ refer to the change in the change in Gibbs free energy, half-maximal inhibitory concentration, half-maximal effective concentration, 80% inhibitory concentration, 50% inhibitory dose and protein-protein dissociation constant respectively.

| Dataset | Raw Datapoints | Mutations | Data Type | Numerical Value |
| --- | --- | --- | --- | --- |
| AB-Bind | 1 101 | Available | Sequences | $\Delta\Delta G$ |
| AB-Cov | 1 964 | Available | Structures | IC50, EC50 |
| CATNAP | 129 686 | Available | Names only | IC50, IC80<br>ID50 |
| SAbDab | 1 327 | Available | Structures | $\Delta\Delta G$<br>Affinity |
| SKEMPI | 7 086 | Available | Structures | Affinity |
| AlphaSeq | 1 259 700 | N/A | Sequences | $K_d$ |

**Fig. 4.**
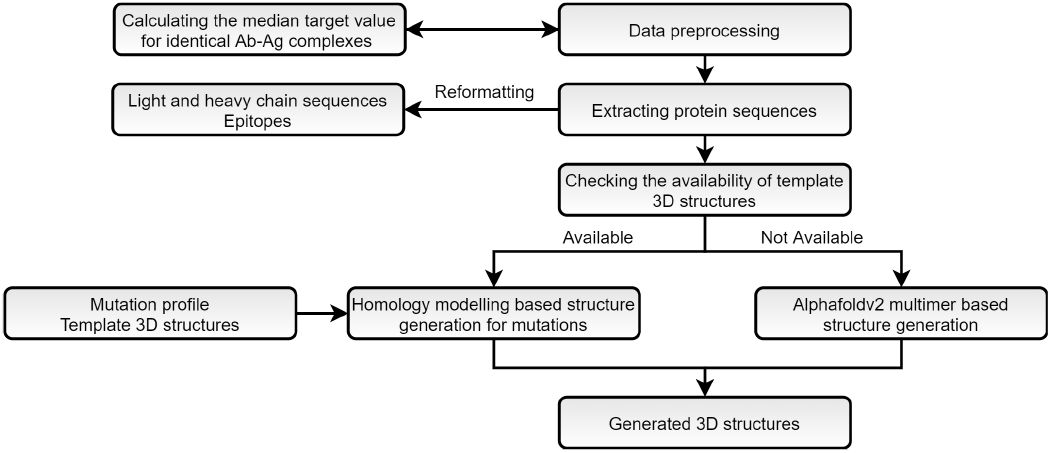
The generalized flowchart describing the steps followed to curate the mentioned datasets.

### B. Homology Modelling

Homology modeling is a reasonably accurate comparative protein structure modeling technique in which an atomic-resolution model of the target protein is constructed using its amino-acid sequence and an experimental 3D structure of a related homologous protein (31). We utilized a homology modeling-based pipeline when the template sequence (original amino acid sequence) and its 3D structure (PDB format) were available along with the mutation profile. First, we created the mutated protein sequence based on the mutation profile provided for the template sequence using a package called MDAnalysis (32). Homology Modelling was then employed to generate the 3D structure (in PDB format) of a mutated sequence (i.e. target sequence) using the template PDB structure. For the sequence alignment task between the target sequence and template sequence, a Python package called Bio-python (33) was used instead of the default sequence alignment function available in the MODELLER for better results.

The first stage in structure derivation was backbone modeling, which is responsible for estimating the positions of the amino group, *α*-Carbon atom, and the Carboxyl group of each amino acid in the polypeptide sequence. Backbone modeling was followed by loop modeling, which performs necessary conformational adjustments to the modeled backbone. Finally, side chains were modeled. To alleviate steric collisions, the estimated relative atomic locations were finetuned to minimize the potential energy of the conformation of the protein. This was done during the model optimization step. At last, the optimized model was evaluated. These steps were conducted using ‘MODELLER’ (34) freeware program.

### C. AlphaFold-V2

AlphaFold-V2 multimer model (35) is a state-of-the-art model with atomistic level accuracy for protein structure prediction from amino acid sequences, which is proven to be useful even in the absence of a homologous structure, as shown by their superior performance at the 14^*th*^ Community Wide Experiment on the Critical Assessment of Techniques for Protein Structure Prediction.

In our implementation for generating the 3D structures of proteins where the template structures were absent, we followed the pipeline from ColabFold (36), which runs inference using the AlphaFold-V2 along with an accelerated combination of MMseqs2-based homology search where MM-seqs2 (37) refers to Many-against-Many-searching which is widely used to search and cluster sequences.

### D. Evaluation Metrics

Throughout the experiments, the following metrics were utilized to evaluate the performances and compare the results between the models.

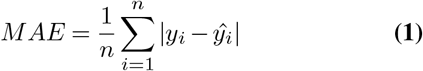

where the evaluation parameter: mean absolute error is denoted as *MAE. y*_*i*_ and ŷ refer to the true value and predicted value, respectively.

The mean squared error (*MSE*) was mainly used as a loss function and on certain occasions, it was also considered as an evaluation metric to further validate the results.

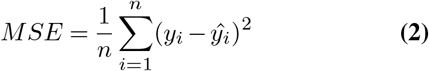

Further, to evaluate the correlation between the true values and predicted values from our models, concerning linearity and ranks, Pearson correlation coefficient (*r*) and Spearman’s correlation coefficient (*ρ*) were utilized respectively.

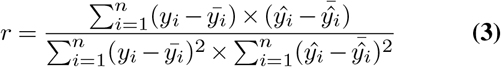

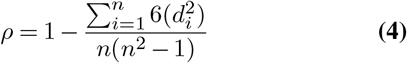

In Equation 4, *d*_*i*_ refers to the difference between the two ranks of each true and predicted value pair.

### E. Sequence-based Model

The input protein files were processed to obtain the FASTA sequences. Since deep learning models require a numerical input we encode the complete FASTA sequence (i.e., to be specific, not just complementarity-determining regions) using a numerical scheme such as one-hot and VHSE8. Suppose the encoded antibody and antigen sequences are of the dimensions *s*_1_ *× d* and *s*_2_*×d*, respectively. Initially, each d-dimensional vector was projected onto another d-dimensional embedding space through a dense layer. Then, the projected matrix was passed through separate, standard attention blocks whose component layers were multi-head attention (self), dropout, layer norm, and dense layers. The equation for the attention mechanism is given by, ∀*i* = 1, 2, 3, …, *s*

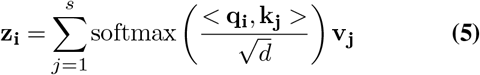

where **z**_**i**_ ∈ ℝ^*d*^ is the output vector after the attention layer, *s* is the sequence length (antibody pathway, *s* = *s*_1_ and antigen pathway, *s* = *s*_2_), *d* is the embedding vector dimension. Furthermore, *<*. *>* indicates the Euclidean inner product. Moreover, **q**_**i**_ = *W*_*q*_**x**_**i**_, **k**_**i**_ = *W*_*k*_**x**_**i**_ and **v**_**i**_ = *W*_*v*_**x**_**i**_ with **x**_**i**_ ∈ ℝ^*d*^ being the input vector and *W*_*q*_, *W*_*k*_, *W*_*v*_ ∈ ℝ^*d×d*^. The soft-max operator is defined as,

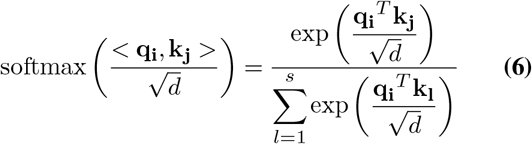

Along with self-attention/multi-head attention layers, cross-attention layers were also implemented to pass information from the antibody pathway to the antigen pathway and vice versa. As shown in Fig. 5, we included two pathways for cross-attention to mimic the hierarchical information sharing. The cross-attention layer has a similar equation as in the case of self-attention with a subtle difference in the limits of summation. Accordingly, for cross-attention, we can provide the following equation considering the flow of information from the antigen pathway to the antibody pathway. ∴ ∀*i* = 1, 2, 3, …, *s*_1_,

**Fig. 5.**
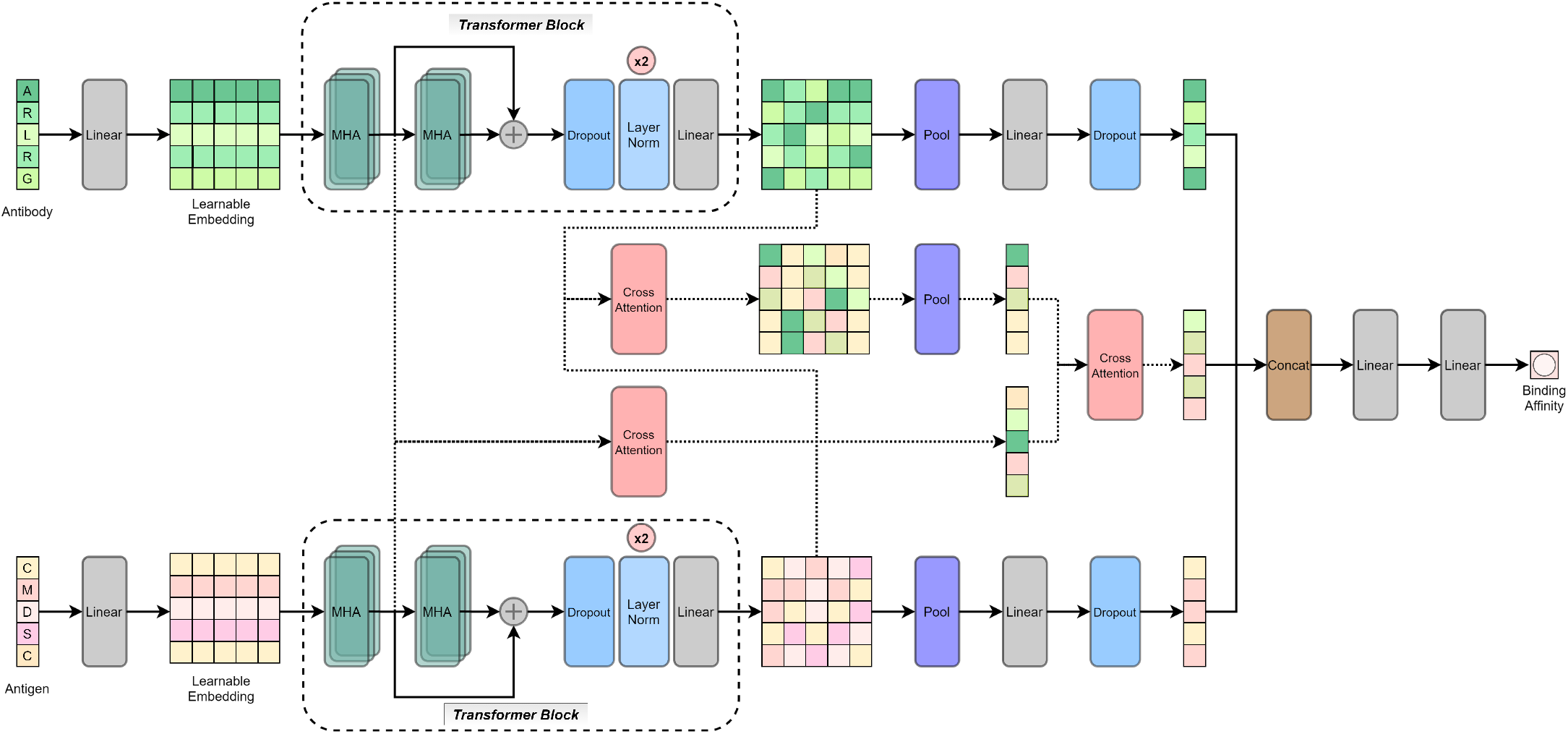
The network architecture for the sequence-based model. *MHA refers to Multi-head Attention and other layers in the diagram hold conventional meanings as used in deep learning.

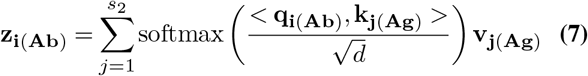

where **z**_**i**(**Ab**)_ ∈ ℝ^*d*^ is the output antibody vector after the cross-attention layer. Moreover, **q**_**i**(**Ab**)_ = *W*_*q*_**x**_**i**(**Ab**)_, **k**_**j**(**Ag**)_ = *W*_*k*_**x**_**i**(**Ag**)_ and **v**_**j**(**Ag**)_ = *W*_*v*_ **x**_**i**(**Ag**)_ with **x**_**i**(**Ab**)_, **x**_**i**(**Ag**)_ ∈ ℝ^*d*^ being the input antibody vector and antigen vector, respectively and *W*_*q*_, *W*_*k*_, *W*_*v*_ ∈ ℝ^*d×d*^. Other symbols hold the same meaning as in the self-attention layer. A similar equation can be provided for the flow of information from the antibody pathway to the antigen pathway by interchanging ‘Ab’ (which refers to antibody) and ‘Ag’ (which refers to antigen) subscripts in Equation 7 and changing the upper limit of summation to *s*_1_ instead of *s*_2_.

The outputs from the antibody pathway, antigen pathway, and antibody-antigen pathways were then concatenated into a single vector and further processed through two dense layers to get the output.

### F. Structure-based Model

Due to the limited information on binding sites for many data-points, here we utilized individual structure details of antibodies and antigens rather than the antibody-antigen complex structure details. In our implementation, we represented both antibody and antigen molecules as graphs 𝒢= (𝒱, ℰ) that use atoms as nodes with their respective 3D coordinates denoted as *X* ∈ ℝ^3*×n*^, and initial atomic features *F* ∈ ℝ^4*×n*^. Edges include all atom pairs within a distance cutoff of 5Å. Initial atomic features include atomic number, implicit valance, charge list, and the degree of the atom. Edge features include the bond strength which we calculate as the reciprocal of bond distance.

The structure model architecture comprises 2 parallel branches that simultaneously process antibody and antigen graph representations, as depicted by Fig. 6. Each branch consists of 4 graph convolutional layers (38), followed by 4 graph attention layers (39) and a graph average pooling layer. To aggregate the features of neighboring atoms, we passed the graph representations of antibody and antigen molecules through a stack of graph convolutional layers. The core property of the graph convolutional layer is that it takes the weighted average of neighbours’ node and edge features, including itself. A single graph convolutional layer, followed by a ReLu non-linear layer, is denoted as follows.

**Fig. 6.**
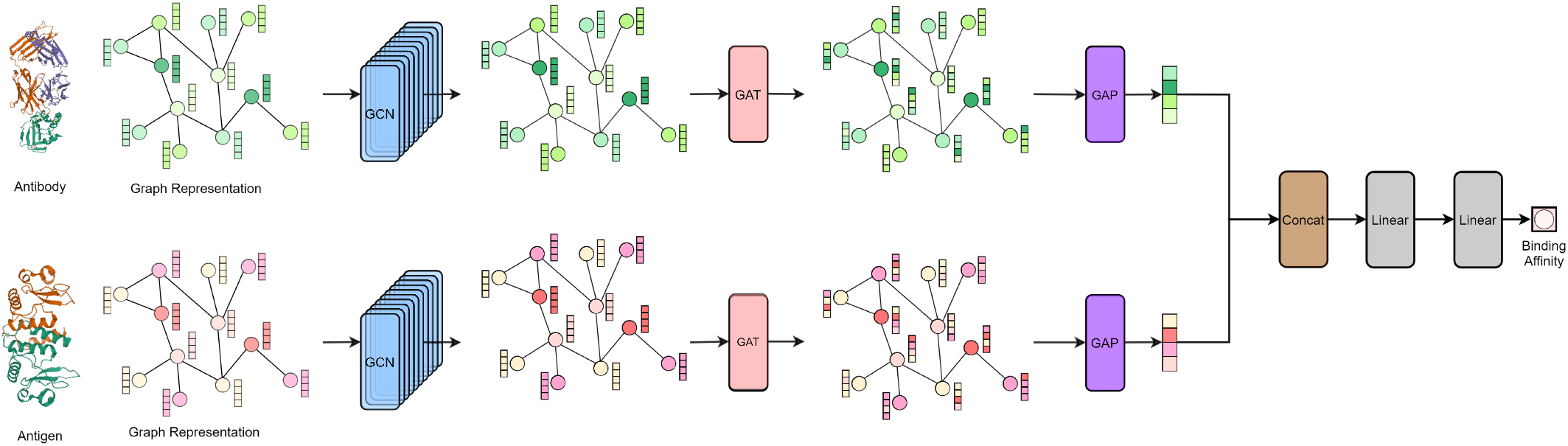
The network architecture for the structure-based model. *Here, GCN, GAT, and GAP refer to Graph Convolution, Graph Attention, and Graph Average Pooling, respectively.

**Fig. 7.**
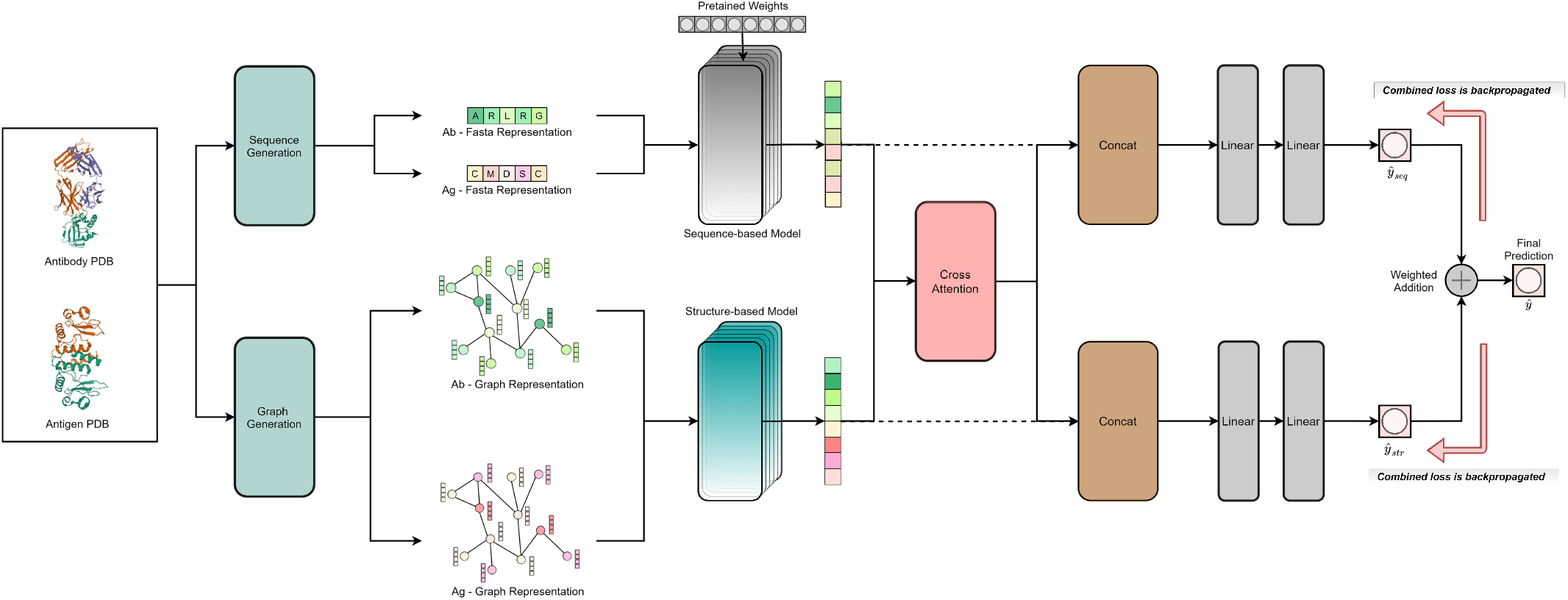
The network architecture for the combined model.

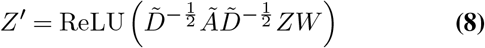

where *Z* ∈ ℝ^8*×n*^ denotes the input feature matrix that includes coordinate values, node, and edge features of n nodes. Here, *A* is the adjacency matrix, Ã = *A* + *I*_*N*_ and 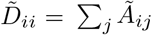. Moreover, *W* is the weight matrix and *I*_*N*_ is the identity matrix. Following the stack of graph convolutional layers, a stack of 4 graph attention layers was applied to identify the atoms that should be given more priority. The graph attention operator works as follows.

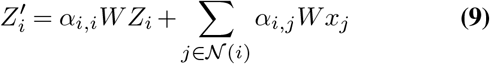

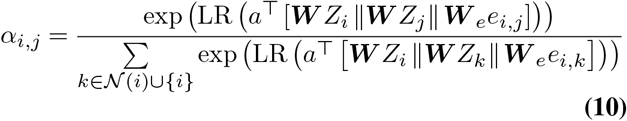

where *Z*_*i*_ is the input feature vector of i th node, *W* is the weight matrix and LR denotes the LeakyReLU non-linear function. Afterwards, we fed the graph attention layer output to a graph average pooling layer to reduce the spatial dimension. Subsequently, embeddings of the antibody and antigen obtained via each pathway were concatenated and processed further through 2 linear (dense) layers to retrieve the binding affinity prediction from the structure model.

### G. Combined Model

With the expectation of capturing evolutionary details through protein amino acid sequences and atomistic details via protein 3D structures to enhance the prediction performance, the final combined model constitutes the aforementioned sequence and structure models. We applied cross-attention to propagate complementary information between the two domains. As indicated by Fig. 7, the cross-attention output was concatenated separately with the embeddings from each constituent model and passed through linear (dense) layers to obtain two intermediate binding affinity predictions (ŷ_*seq*_ and ŷ_*str*_). The final binding affinity prediction (ŷ) is a weighted average between those two predictions, and the weights were decided based on the conducted ablation study.

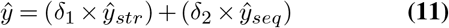

Here, we used a weighted average function with hyperparameters; *δ*_1_ and *δ*_2_. Through ablations, *δ*_1_ and *δ*_2_ were selected to be 0.45 and 0.55, respectively.

During the training phase of the combined model, for a given ground truth value (*y*), the following combined loss function (ℒ_*total*_) was calculated, and the error was backpropagated, updating the weights in the constituent sequence and structure models, simultaneously.

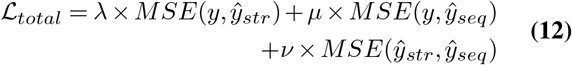

where *λ, µ* and *ν* are hyperparameters selected through ablation studies. The intermediate binding affinity predictions from the sequence and structure models are denoted by ŷ_*seq*_ and ŷ_*str*_, respectively.

## Supporting information

High-level Overview

## Appendix

### A. Ablations on Sequence Encoding Schemes

As discussed in the section D, it is required to convert the letter sequences of protein amino acids into a numerical format before feeding into the sequence-based models. Accordingly, two encoding schemes were utilized for this task, namely, one-hot encoding and VHSE8 encoding. As indicated in Table 7, the VHSE8 encoding scheme was utilized in the model implementations (including sequence-based models and the combined models) due to its superior performance over the one-hot encoding scheme. We believe that the chemical and physical properties of the amino acids embedded within the VHSE8 encoding scheme are the reason for the better performance compared to one-hot encoding, which is a sparse encoding scheme.

**Table 7.** Results comparison between different encoding schemes. Here, the P2PXML-Seq dataset is utilized and the model is a parallel multi-layer perception model.

| Encoding scheme | MAE |
| --- | --- |
| One-hot | 0.9757 |
| VHSE8 | **0.9682** |

As discussed previously under the Results section, the protein laguage embeddings from ProtT5-XL-BFD (25) were also utilized to numerically encode the protein amino acid sequences. However, it was shown in Table 3 that the performance of such an embedding representation is inferior to the VHSE8 encoding scheme in this context.

### B. Ablations on Model/Training Parameter Selections

It is important to run extensive ablations for different combinations of hyperparameters as we were trying to heuristically determine the parameters that would maximize the performance of the model. Accordingly, as per Table 8, it is evident that the best MAE is given by a learning rate of 0.0001, along with the ADAM optimizer and a dropout of 0.05. The set of hyperparameters that exhibit the best MSE is almost similar to those of the best MAE except for the dropout, which is 0.10 in the case of the best MSE. However, under the hyper-parameters that produced the best MSE, we obtain an MAE that is relatively worse than the best MAE. Moreover, note that the MSE corresponding to the best MAE is quite close to the best MSE. Accordingly, it was decided to use the set of hyperparameters that produced the best MAE value.

**Table 8.** Optimal hyperparameter determination through ablations by exposing the proposed sequence model to the AlphaSeq dataset. *LR, D/O, EV refer to Learning Rate, Dropout and Embedding Vector, respectively.

| LR | Optimizer | D/O | EV | MAE | MSE |
| --- | --- | --- | --- | --- | --- |
| 0.001 | ADAM | 0.05 | ✗ | 0.179275 | 0.049301 |
|  |  | 0.1 | ✗ | 0.335331 | 0.116954 |
|  |  | 0.2 | ✗ | 0.130520 | 0.039792 |
| 0.0001 | ADAM | 0.05 | ✗ | **0.054212** | 0.036044 |
|  |  | 0.1 | ✗ | 0.088051 | 0.036068 |
|  |  | 0.2 | ✗ | 0.058286 | 0.035924 |
|  |  | 0.05 | ✓ | 0.103310 | 0.036906 |
|  |  | 0.1 | ✓ | 0.066851 | **0.035740** |
|  |  | 0.2 | ✓ | 0.062289 | 0.035802 |
|  | ADA-DELTA | 0.05 | ✗ | 0.089726 | 0.036105 |
|  |  | 0.1 | ✗ | 0.091582 | 0.036193 |
|  |  | 0.2 | ✗ | 0.094721 | 0.037684 |

### C. Ablations on Graph Node and Edge Features

One of the most crucial steps in geometric deep learning, specifically graph-based deep learning, is the initial graph representation, as it would directly impact the subsequent node and edge level aggregations and information flow.

Therefore, an extensive ablation study was performed to find an optimal set of node and edge features for the graph representation of proteins and thereby strengthen the chemistry-based intuition of the final model. The results under two datasets, namely, Ab-CoV and Ab-Bind were used to compare different combinations. As per the results presented in Table 9, the following set of node features: x-coordinate, y-coordinate, z-coordinate, Atomic Number, Implicit Valence, Charge List, Degree, and edge features: Bond Strength and Atomic Distance were selected to be used in the remaining experiments associated with the structure-based model and the combined model.

**Table 9.**
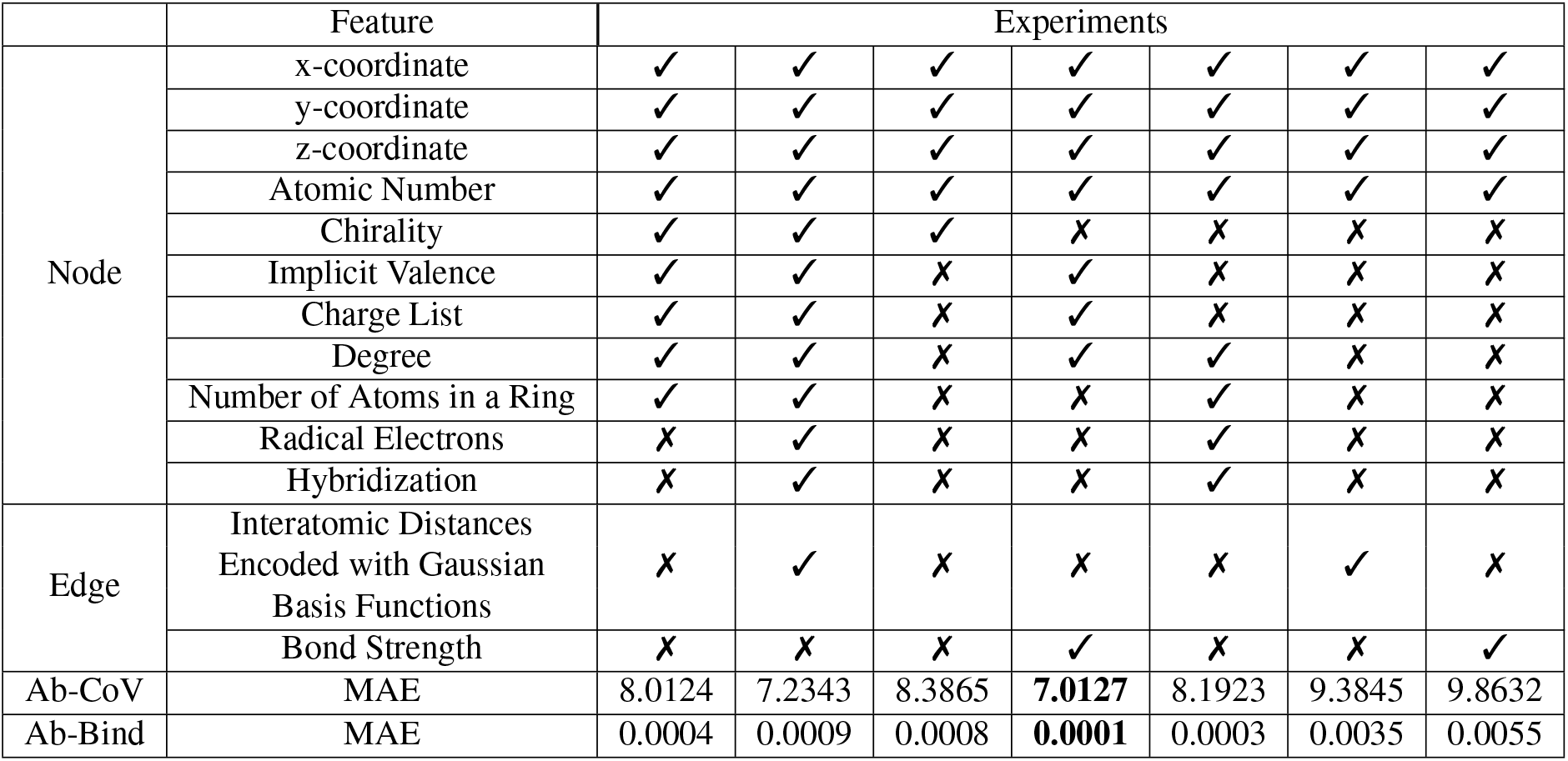
Evaluation of node and edge features using the protein structures generated (using AlphaFold-V2 multimer model) for the protein sequences in Ab-Cov and AB-Bind datasets

### D. Timing Analysis

One of the most critical challenges in traditional wet-lab-based or molecular dynamics-based experiments for antibody-antigen binding affinity estimation is the time complexity. However, through deep learning-based approaches, it is possible to obtain an accurate prediction for the binding affinity within minutes or even seconds. Accordingly, our Combined-V2 model is able to predict the binding affinity of a given antibody-antigen pair within 1 minute, provided a graphic processing unit (GPU) and within 3 minutes on a CPU as shown in Table 10.

**Table 10.**
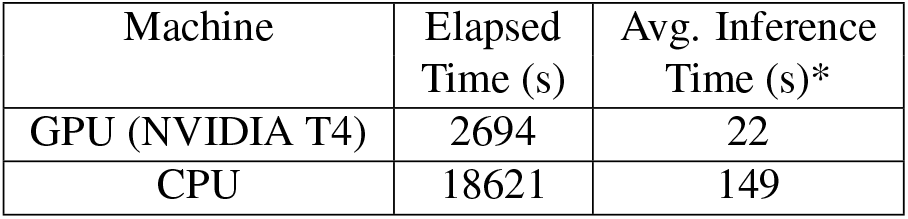
Timing analysis for the Combined-V2 model. *Here, the elapsed time refers to the time taken to run inference on a random sample of 125 antibody-antigen pairs from the P2PXML-PDB dataset. The times are expressed to the nearest second.

**Table 11.** The input format for each model hosted on the website. The units of the output are the predicted binding affinity in *µg/ml*. The validity of the inputs is checked before feeding to our models through rule-based conditioning.

| Model | Input format |
| --- | --- |
| Combined-V2 | PDB files |
| Final sequence-based model | Text sequences |
| Final structure-based model | PDB files |

### E. Website, Project Page and Code Availability

The web platform is developed as a community access tool where anyone interested can obtain the outputs from our Combined-V2, final sequence-based, and structure-based models for their input protein sequences and/or PDB files. The input format for each model on the website is as follows:

The website is accessible through this link: https://p2pxml.azurewebsites.net/ while the multimedia materials, including the curated datasets, demonstration videos and the codes, will be available on the project page: https://drug-discovery-entc.github.io/p2pxml/.

## ACKNOWLEDGEMENTS

We are grateful to Dr. Ranga Rodrigo, Former Head of the Department of Electronic and Telecommunication Engineering for allocating us GPUs which significantly accelerated the protein structure generation process and training the deep learning models. We would also like to extend our gratitude to other Senior Lecturers at the Department of Electronic and Telecommunication Engineering, University of Moratuwa for the valuable feedback given during the project implementations.

